# Antibiotic resistance via bacterial cell shape-shifting

**DOI:** 10.1101/2021.10.16.464635

**Authors:** Nikola Ojkic, Diana Serbanescu, Shiladitya Banerjee

## Abstract

Bacteria have evolved to develop multiple strategies for antibiotic resistance by effectively reducing intra-cellular antibiotic concentrations or antibiotic binding affinities, but the role of cell morphology on antibiotic resistance remains poorly characterized. By analyzing cell morphological data of different bacterial species under antibiotic stress, we find that bacterial cells robustly reduce surface-to-volume ratio in response to most types of antibiotics. Using quantitative modelling we show that by reducing surface-to-volume ratio, bacteria can effectively reduce intracellular antibiotic concentration by decreasing antibiotic influx. The model predicts that bacteria can increase surface-to-volume ratio to promote antibiotic dilution if efflux pump activity is reduced, in agreement with data on membrane-transport inhibitors. Using the particular example of ribosome-targeting antibiotics, we present a systems-level model for the regulation of cell shape under antibiotic stress, and discuss feedback mechanisms that bacteria can harness to increase their fitness in the presence of antibiotics.

## Introduction

Antibiotic resistance is one of the major threats to human society. It has been estimated that each year 700,000 people die as a consequence of infections caused by resistant bacteria, prompting urgent response in order to prevent devastating global effects within generations [1]. To understand the mechanisms of antibiotic resistance we need to better understand how antibiotics physically penetrate bacterial cells, how antibiotics bind to their targets, what damage antibiotics cause to bacterial physiology and ultimately how this damage leads to cell death [2, 3]. To become antibiotic resistant, bacteria have developed multiple strategies. Resistance is commonly attained by via a reduction in the intracellular concentration of the antibiotic or by reducing antibiotic binding affinities to their specific intracellular targets (Figure 1A) [4]. Varioius different pathways to antibiotic resistance have been described [5] including decrease in antibiotic influx by reduction in porin expression [6], modulation of membrane lipid composition [7], induction of horizontal gene transfer, increase in antibiotic efflux by increasing efflux pump expression [8], SOS response [9], and direct inactivation of antibiotics [10]. However, the role of cell size, shape and growth physiology on antibiotic resistance remains poorly understood.

**Figure 1.**
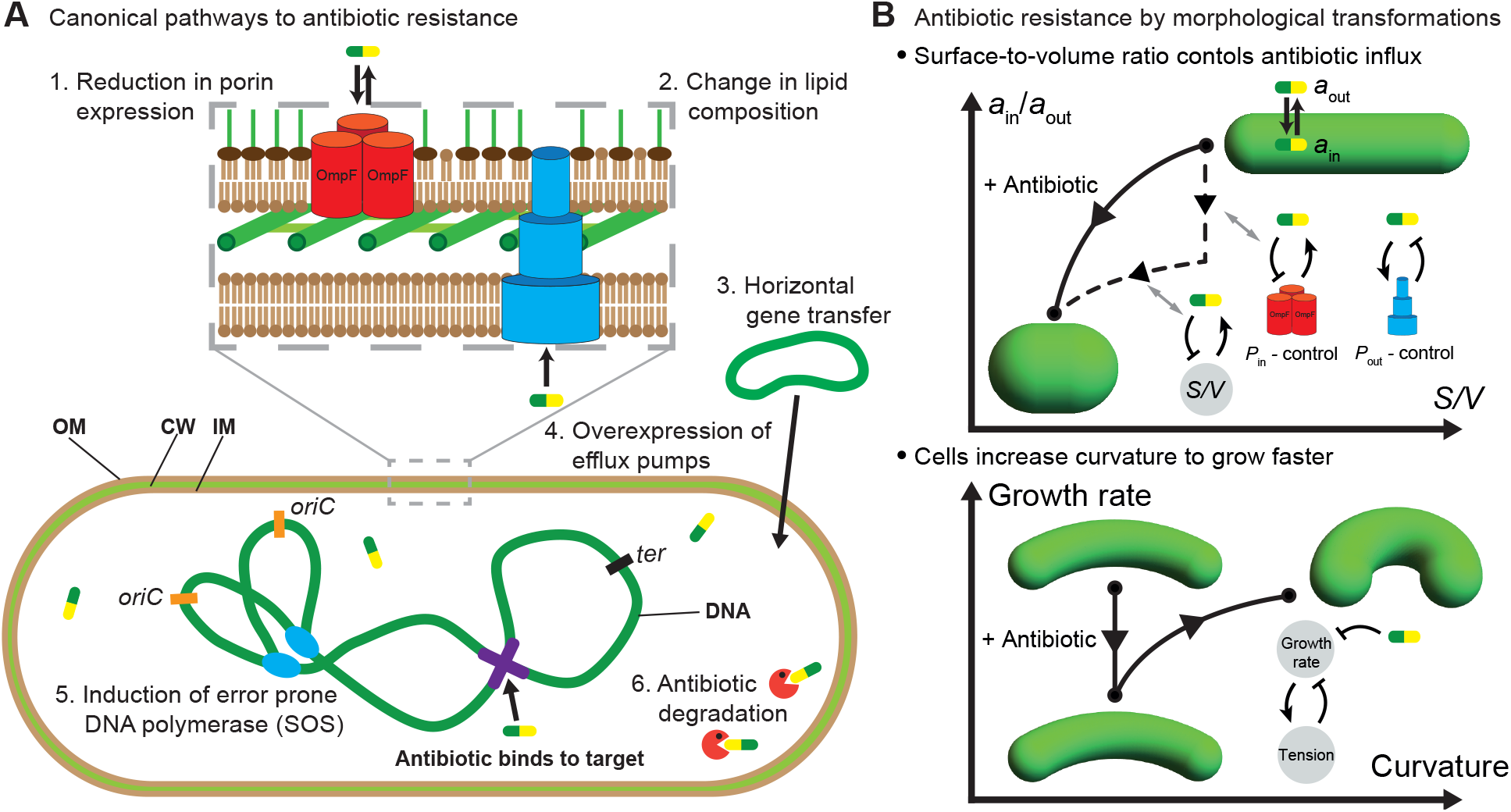
Mechanisms of antibiotic resistance at the single-cell level. (A) Canonical mechanisms of antibiotic resistance result in reduced intracellular antibiotic concentration or reduced antibiotic binding affinities to their targets. Six pathways are shown: (1) Reduction in porin expression. Trimer of passive transporter porin OmpF shown in orange. (2) Lipid composition affects antibiotic translocation across the membrane. (3) Acquiring resistant genes through horizontal gene transfer. (4) Over expression of efflux pumps depletes intercellular antibiotics. Multi drug efflux pump AcrAB-TolC shown in blue. (5) During SOS response bacteria express error prone DNA polymerases that increase random mutation rate. (6) Antibiotics are neutralised by specific proteins shown as red pacmans. Example of antibiotic binding to the target (DNA gyrase) shown in purple [11]. OM - outer membrane, IM - inner membrane, CW - cell wall. (B) Emerging pathways of antibiotic resistance via cell shape transformation. (Top) Reduction in antibiotic concentration inside of the cell through surface-to-volume changes. (Bottom) By altering cell geometry and cell envelope mechanics bacteria promote faster growth.

Recent studies have shown that bacteria undergo a wide variety of cell morphological changes in response to antibiotics [12–18]. These morphological changes (Fig. 1B) commonly occur via changes in cell size, surface-to-volume ratio or curvature [13, 15, 18]. For instance, Gram-negative *E. coli, C. crescentus* and Gram-positive *L. monocytogenes* decrease their surface-to-volume ratio (*S/V*) upon treatment with ribosome-inhibitory and cell-wall targeting antibiotics [15]. It has also been shown that the Gram-negative human pathogen *Pseudomonas aeruginosa* make a transition from rod-shaped cells to spherical cells upon treatment with *β*-lactams [14]. However, it is not clear if these shape changes represent a passive physio-logical response to biochemical changes caused by the antibiotic, or if these are active shape changes that promote bacterial fitness for surviving antibiotic exposure. While the role of cell shape on bacterial growth, nutrient uptake and motility have been hypothesized [19], the effect of cell shape on antibiotic resistance remains largely unknown. It remains unclear whether different classes of antibiotics that target distinct cellular components and physiological processes would increase or decrease *S/V*.

In this article, we use quantitative modeling and data analysis to propose that cell shape-shifting via changes in *S/V* promote antibiotic resistance in bacteria. In particular, by developing a mathematical model for antibiotic transport and binding coupled to bacterial cell shape and growth, we show that by changing *S/V* of the cell, bacteria can effectively dilute intracellular antibiotic concentration by decreasing antibiotic influx (Fig. 1B). The model allows us to predict the quantitative range for antibiotic dilution via reduction in *S/V* . The model also explains that bacteria can increase *S/V* to reduce intracellular antibiotic concentration if the activity of the efflux pumps are reduced, in agreement with data on membrane-transport inhibitors. To understand how antibiotics induce cell shape transformations, we develop a systems-level model for cell shape regulation using the particular example of ribosome-targeting antibiotics for which the biochemical pathways are quantitatively characterized. In addition to providing a systems-level model for the antibiotic adaptation by shape changes, we discuss other adaptation mechanisms in which bacteria alter cell-envelope mechanics to undergo shape changes, which in turn restore growth rate to pre-antibiotic values. We conclude by discussing additional mechanisms of countering antibiotics via regulating metabolic pathways, membrane porin and efflux pump expressions that may act in synergy with shape-induced resistance.

### Antibiotic-induced cell shape changes in rod-shaped bacteria

To understand the effect of antibiotics on bacterial cell shape, we first analysed the morphological data for *E. coli* cells treated with a total of 42 antibiotics belonging to 5 different categories based on their binding targets (Fig. 2A) [13]. Surprisingly, all antibiotics decrease *S/V* except membrane- and membrane transport-targeting antibiotics that increase *S/V* . Similarly, decrease in *S/V* was previously observed in cells treated with cell-wall targeting antibiotics (A22, mecillinam, fosfomycin, cephalexin) [22], and increase in *S/V* for the membrane targeting antibiotic cerulenin [23]. We find that the Gram-negative *A. baumannii* also decreases *S/V* for most antibiotics, including the membrane targeting antibiotic triclosan (Fig. 2B). For Gram-positive *B. subtilis* with thick, less plastic cell-envelope [24, 25], *S/V* decreases for all groups of antibiotics (Fig. 2C). To understand the implications of these data, we note that *S/V* is one of the key physical parameters that regulates nutrient influx and waste efflux [19], as well as the influx/efflux of antibiotics. To quantitatively understand the role of *S/V* in regulating antibiotic flux across the cell membrane, we developed a mathematical model of antibiotic transport into a rod-shaped bacterial cell, with binding/unbinding interactions with its specific target.

**Figure 2.**
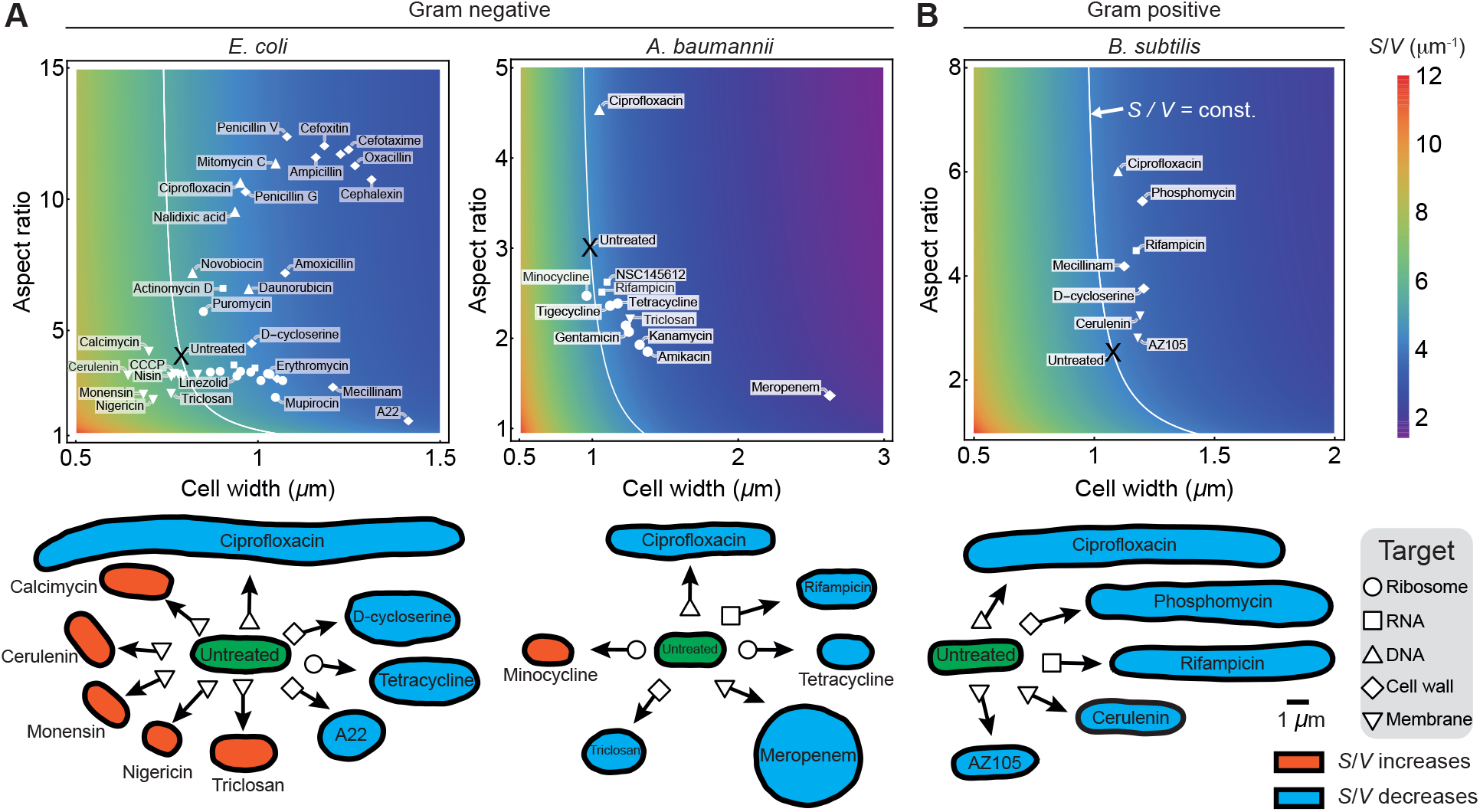
Changes in cell shape and surface-to-volume ratio of rod-shaped bacteria under different antibiotics. Top row: Heatmap of cell surface-to-volume ratio (*S/V*) as a function of cell width and aspect ratio, overlayed with experimental data for cell shape under antibiotic treatments targeting different cellular components: ribosomes, RNA, DNA, cell wall, and membranes. White lines represent constant *S/V* corresponding to untreated cells. Bottom row: Typical cell contours for morphological response to antibiotic treatments. *S/V* increase shown in red, decrease in blue, and untreated cells in green. Data are taken from refs [13, 20, 21]. (A) *S/V* for Gram-negative *E. coli* and *A. baumannii* as a function of cell width and cell aspect ratio. *E. coli* decrease *S/V* for all antibiotics apart from membrane targeting ones for which *S/V* increases. *A. baumannii* decreases *S/V* for all antibiotics apart for ribosome targeting Minocycline for which *S/V* slightly increases. (B) *S/V* for gram-positive *B. subtilis* decreases for all antibiotics.

### Cellular surface-to-volume ratio regulates intracellular antibiotic concentration

When antibiotics are passively translocated into the cell thorough membrane porins or lipids (Fig. 3A), antibiotic transport is diffusion limited and the flux is given by Fick’s law [26, 27]. Dynamics of intracellular antibiotic concentration, *a*_in_, and the substrate concentration, *x*, are given by:

**Figure 3.**
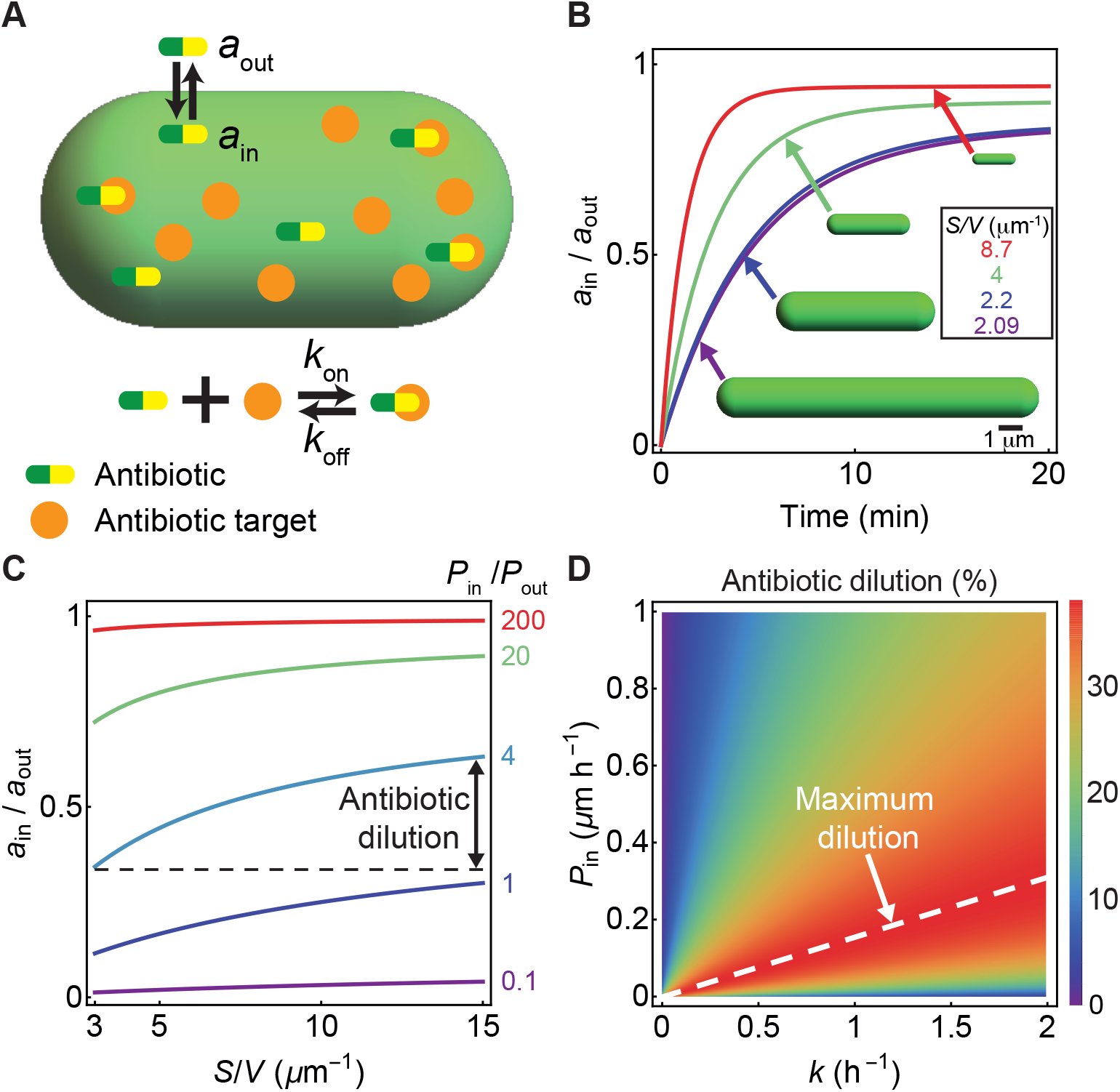
Cell shape-dependent dynamics of antibiotic concentration in a bacterial cell. (A) Schematic of the model for antibiotic uptake and substrate binding/unbinding inside a bacterial cell. (B) Relative antibiotic con-centration inside a cell vs time for different values of *S/V*, with all other parameters kept fixed: *P*_in_ = 5 *μ*m h^*−*1^, *P*_out_ = 0.1 *μ*m h^*−*1^, *k*_on_ = 1 *μ*M^*−*1^ h^*−*1^, *k*_off_ = 0 h^*−*1^, *k*_*x*_ = 1 *μ*M h^*−*1^, *k* = 1 h^*−*1^. (C) Steady-state antibiotic concen-tration, normalized by *a*_out_, vs *S/V* for different values of *P*_in_*/P*_out_. For the cases *P*_in_ ≫ *P*_out_ ≪ or *P*_in_ *P*_out_ antibiotic dilution due to *S/V* changes is negligible. (D) Heat map of percent antibiotic dilution (100*δ*) for different values of *k* and *P*_in_, setting *P*_out_ = 0, (*S/V*)_min_ = 3 *μ*m^*−*1^, (*S/V*)_max_ = 15 *μ*m^*−*1^. Maximum antibiotic dilution, predicted by Eq. (4), is shown by the white dashed line.

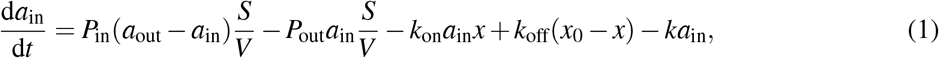

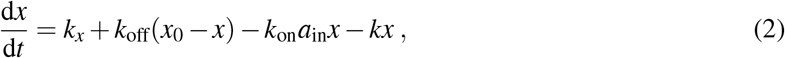

where *a*_in_ is the concentration of antibiotic in the extracellular medium, *x*_0_ is the substrate concentration in the absence of antibiotic, *k*_on_ is the antibiotic binding rate to the substrate, *k*_off_ is the antibiotic unbinding rate, *P*_in_ and *P*_out_ are the membrane permeability coefficients in the inward and outward directions, respectively. The last term on the right hand side of Eqs. (1) and (2) represents the dilution of the antibiotic and the substrate due to cell growth, where *k* is the bacterial growth rate. Numerical solution of the above system of equations for different values of *S/V* predicts the time-evolution of the intracellular antibiotic concentration (Fig. 3B), where the steady-state concentration of the antibiotic decreases with decreasing *S/V* (Fig. 3C). Since all antibiotics, apart from membrane-targeting ones, decrease *S/V* in bacteria, these results point towards an adaptive strategy to reduce intracellular antibiotic concentrations via geometric changes. To conceptually understand these numerical results, we note that flux balance is reached at steady-state:

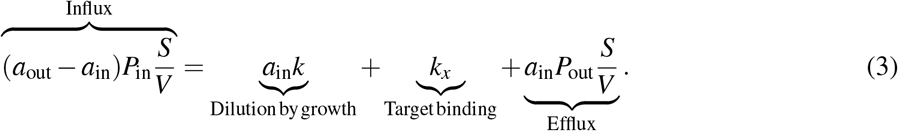

Therefore, the strategy to decrease *S/V* upon antibiotic treatment (Fig. 2), results in a reduction in the antibiotic influx term on the left hand side of Eq. (3). In the case when efflux pumps are effective (*P*_out_ *>* 0) *S/V* reduction also leads to an overall decrease in antibiotic efflux. Conversely, when *E. coli* cells are exposed to membrane-targeting and membrane transport-targeting antibiotics, bacterial surface area decreases whereas *S/V* increases. The decrease in surface area is expected, however benefits of *S/V* increase could be interpreted as follows. For the antibiotics that disrupt proton motive force (calcimycin, CCCP, DNP, monensin, nigericin) and therefore efflux pumps efficiency, bacteria can counterbalance complete reduction in overall efflux efficiency (last term in the Eq. 3) by increasing *S/V* . These results prompt for more experimental data on the role of *S/V* in bacterial morphological adaptation depending on the antibiotic targets.

To quantify the effect of *S/V* reduction on intracellular antibiotic dilution (Fig. 3C), we define the *antibiotic dilution factor δ* ≡ |Δ*a*_in_|*/a*_out_, where Δ*a*_in_ is the change in intracellular antibiotic concentration as *S/V* is varied between a chosen minimum (*S/V*)_min_ and a maximum value (*S/V*)_max_. For a given cell, the maximum dilution factor can be analytically calculated as (Fig. 3D and Supplementary Information):

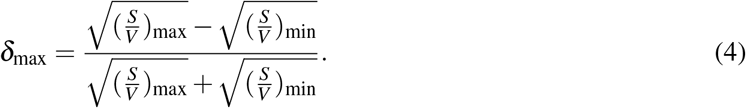

The above equation predicts a maximum *≈* 15% dilution in intracellular antibiotic concentration for cephalexin treated *E. coli*, where (*S/V*)_max_ and (*S/V*)_min_ are taken to be the surface-to-volume ratios for the untreated cell and the antibiotic-treated cell, respectively. Similarly, meropenam treated *A. baumannii* cells undergo a maximum of 22% dilution in intracellular antibiotic concentration. Antibiotic dilution mediated by changes in *S/V* could provide a significant fitness advantage for antibiotics with steep growth-inhibition curves [11, 28]. For spherocylindrical cells of width *w*, Eq. (4) can be simplified as:

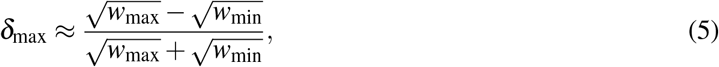

where *w*_min_ and *w*_max_ are the minimum and the maximum width of the cell. Since antibiotic dilution predominantly depends on cell width, a central question is how the width is determined and how fast it is remodelled in response to antibiotic treatments [29–31]. During steady-state growth, cell width is one of the most tightly controlled cellular parameters in both *E. coli* and *B. subtilis* [32], suggesting that bacteria tightly regulate *S/V* at the single-cell level [22]. During steady-state exponential growth in *E. coli*, fluctuations in width are restored within 4-5 generation [17], commensurate with the typical time necessary for bacterial width remodelling in agreement with theoretical predictions [33]. In non-steady conditions, the cell width reaches a new equilibrium only within a few generations after nutrient shifts [34] or antibiotic exposure [15, 18]. Cell width control is achieved by the cell wall synthesis machinery such as MreB, RodZ and penicillin-binding proteins [35], as well as by physical forces, including mechanical stress on the cell envelope and osmotic pressure [36–38]. Understanding the mechanisms of cell shape regulation therefore requires an integrative approach, combining cellular mechanics and biochemical regulation.

### Mechanisms of cell shape changes by antibiotic action

The mechanisms by which antibiotics induce cell shape changes are specific to the type of antibiotic treatment [13]. While DNA-targeting antibiotics induce SOS response that in turn inhibits cell division causing cell filamentation [11, 13], cell-wall targeting antibiotics such as *β* -lactams induce various cell shapes and sizes (Fig. 2) and cell bulging [12]. Ribosome-targeting antibiotics on the other hand induce a less complex variety of shape changes [15, 18, 39]. Here we specifically focus on the case of ribosome-targeting antibiotics, for which the biochemical reactions are well characterized [40], in order to elucidate the dynamic coupling between cell shape, growth rate and antibiotic concentration.

One of the most commonly used ribosome-targeting antibiotics is chloramphenicol (CHL), which inhibits bacterial translation. When actively translating ribosomes are blocked by CHL, cells synthesize new ribosomes in excess to counterbalance their inactivation [41]. This increase in ribosome abundance requires bacteria to strategically focus their ribosomal resources towards growth rather than division, causing an increase in cell volume [42]. Interestingly, when *E. coli* cells are exposed to nutrient or CHL perturbations bacterial cell aspect ratio remains constant *≈* 4 [17], implying a simple scaling relation between surface area and volume, *S* = 2*πV* ^2*/*3^, yielding *S/V* = 2*πV*^*−*1*/*3^ [17]. Therefore, if the bacteria increase volume upon CHL treatment, due to excess ribosome synthesis, *S/V* would then decrease (Fig. 4A-B). More specifically in *E. coli* cells, *S/V* = 2*π*(*k*_*F*_ */k*)^1*/*3^ (Supplementary material), where *k* is the growth rate and *k*_*F*_ is the volume-specific rate of synthesis of division proteins (Fig. 4A) [42]. Both *k*_*F*_ and *k* are regulated by ribosomes, such that under translation inhibition *k*_*F*_ */k* decreases to favor growth over division. However, depending on the nutrient conditions the ratio *k*_*F*_ */k* sometimes increases. This particularly happens in nutrient-rich medium when cells allocate more resources to division than growth under antibiotic stress, leading to an increase in the surface-to-volume ratio [39, 42].

**Figure 4.**
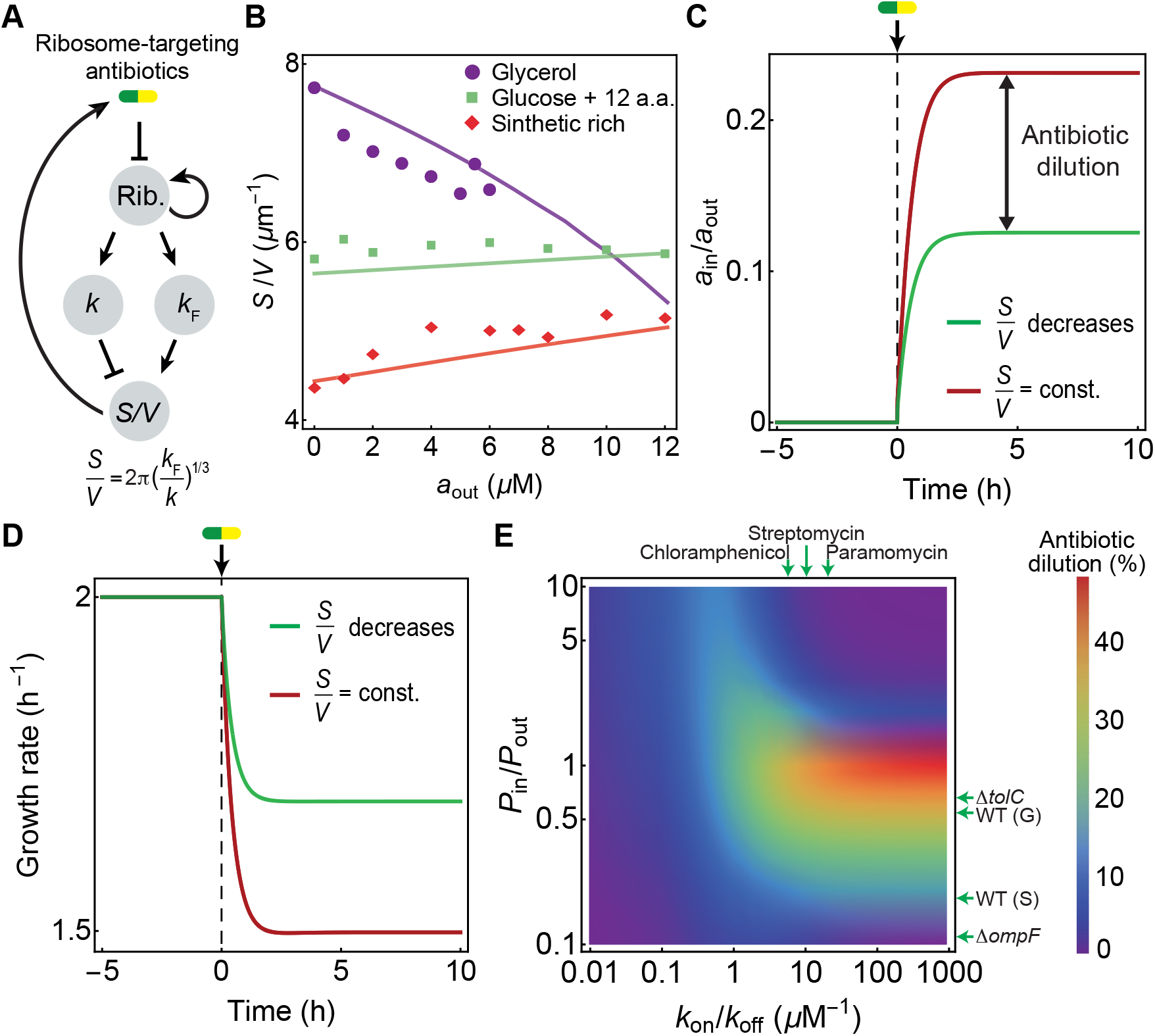
Interplay between cell shape and translation inhibiting antibiotics. (A) Schematic diagram of the feedback pathways connecting ribosomal translation to cell shape and antibiotic transport. Ribosomes promote growth which in turn decreases the surface-to-volume ratio (*S/V*), and an increase the division proteins production rate (*k*_*F*_) increases the surface-to-volume ratio. *S/V* promotes antibiotic influx (Fig. 1B, Eq. 1). (B) *S/V* vs chloramphenicol concentration in *E. coli* [39, 42]. Experimental data for different nutrient conditions are shown as scatter points, simulations are shown as solid lines. (C) Relative antibiotic concentration inside the cell vs time obtained by model simulations for two cases: (i) *S/V* = const. = 5 *μm*^*−*1^ (red), (ii) *S/V* decreases from 5 to 3 *μm*^*−*1^ (green) via the pathway shown in panel (A). Here *a*_out_ = 5 *μ*M and *P*_in_*/P*_out_ = 1. (D) Bacterial growth rate vs time obtained from simulations for two different cases as in panel C. (E) Heat map of antibiotic dilution factor predicted from simulations as functions of membrane permeability ratios (*P*_in_*/P*_out_) and the ratio of antibiotic-ribosome binding and unbinding rates (*k*_on_*/k*_off_). Antibiotic dilution was calculated when bacterial *S/V* was altered from 15 to 3 *μm*^*−*1^, as in Fig. 3C. Membrane permeability ratios shown with horizontal arrows were estimated for growing wild-type, WT (G), and stationary wild type *E. coli* cells, WT (S), OmpF porin deficient (Δ*ompF*) and efflux-pump deficient (Δ*tolC*) cells. Experimentally measured *k*_on_*/k*_off_ values for different ribosome-targeting antibiotics are shown with vertical arrows.

To estimate the amount of antibiotic dilution due to cell shape changes, we simulated a whole-cell model of bacterial growth that evolved the dynamics of cell size, shape and protein synthesis, coupled to the kinetics of antibiotics binding to and unbinding from ribosomes (Supplementary material) [42]. In this model, cell volume grows exponentially at a rate *k*, whereas cell surface area and division proteins are synthesized at rates proportional to cell volume. Using parameters benchmarked for *E. coli* cells, we find that under conditions when *S/V* remains unchanged, intracellular antibiotic concentration is higher and the growth rate is lower compared to the case where *S/V* spontaneously decreased over time (Fig. 4C-D). Our simulations revealed that the maximum antibiotic dilution was obtained for antibiotics with high affinity constants (*k*_*A*_ = *k*_on_*/k*_off_ *>* 1*μ*M^*−*1^) typical for aminoglycosides: Hygromycin B (*k*_*A*_ = 5 *μ*M^*−*1^), Chloramphenicol (*k*_*A*_ = 5.8 *μ*M^*−*1^), Streptomycin (*k*_*A*_ = 10 *μ*M^*−*1^), and Paromomycin (*k*_*A*_ = 19 *μ*M^*−*1^). Depending on the ratio of the membrane permeability constants (*P*_in_*/P*_out_), the antibiotic dilution factor is non-monotonic and reaches a maximum for *P*_in_*/P*_out_ *≈* 1 (Fig. 3C and Fig. 4E). Since Chloramphenicol is predominantly transported inside of the cell by OmpF porins, we estimated the permeability coefficients for fluorescent oflaxacin that is also translocated by OmpF [43]. By analyzing time traces of fluorescent antibiotic accumulation inside of the cell, we estimated that growing *E. coli* cells readily accumulate antibiotics, *P*_in_*/P*_out_ *≈* 0.54, while for stationary cells *P*_in_*/P*_out_ *≈* 0.11. Interestingly, for bacterial cells lacking porins (Δ*ompF*), *P*_in_*/P*_out_ *≈* 0.18 and for bacteria lacking efflux pumps (Δ*tolC*), *P*_in_*/P*_out_ *≈* 0.66 (see Supplementary material). Therefore, these results suggest that the antibiotic dilution by changes in *S/V* could approach the maximum value achieved for *P*_in_ *≈ P*_out_ and large *k*_on_*/k*_off_ (Fig. 4E).

## Conclusion

While rod-shaped bacteria reduce their surface-to-volume ratio upon antibiotic treatment, recent work found that Vibrio bacteria can also alter curvature when exposed to antibiotics [18]. In particular, when *C. crescentus* is treated with sub-MIC concentrations of chloramphenicol, the cell becomes more curved and the cell width increases [18]. Immediately upon CHL treatment cell growth rate decreases, but over *≈* 10 generations the growth rate gradually restores to the pre-antibiotic level. While the cell *S/V* decreases during CHL exposure, the contribution of lowered *S/V* cannot solely explain the almost full growth-rate recovery. This adaptive response via cell shape changes can be explained by a model of negative feedback between cellular growth rate and cell envelope mechanical tension (Fig. 1B) [18]. Translation inhibition by CHL reduces the rate of synthesis cell envelope material, which leads to an initial fast drop in growth rate. The reduced rate of surface area synthesis also reduces the effective tension that works against the compressive bending forces acting on the cell surface. As a result, reduced tension leads to cell surface bending until a new mechanical equilibrium is reached with a higher curvature. Lower cell envelope tension promotes cell-wall synthesis [37], thereby increasing the growth rate to its pre-stimulus value. Future experiments are needed to further test the role of cell curvature and wall tension on the maintenance of growth rate homeostasis. In particular, it will be interesting to test whether mutant Caulobacter cells with straight morphologies are poor at adapting to chloramphenicol, or whether Vibrio bacteria adapt better than straight rod-shaped cells.

A reduction in surface-to-volume ratio not only lowers antibiotic influx, but also leads to a reduced nutrient influx that may in turn lower cellular metabolic activity. In recent work, Lopatkin *et al*. showed that metabolism plays a crucial role in bacterial response to antibiotics such that cells with decreased metabolic activities are more antibiotic-resistant [44]. Metabolic mutations in response to antibiotic exposure suggest adaptive mechanisms in central carbon and energy metabolism. Interestingly, some of the advantageous metabolic mutations that mitigate antibiotic susceptibility have been identified in *>* 3500 clinically relevant pathogenic *E. coli* [44]. These findings point towards a new pathway of antibiotic resistance mediated by mutations in the core metabolic genes. Our findings that bacteria decrease surface-to-volume may act in synergy with the metabolic slowdown to confer stronger antibiotic resistance.

In synergy with shape changes, bacteria can actively regulate antibiotic concentration inside of the cell by controlling porin and efflux-pump expressions (Fig. 1B) [45, 46]. Cell-wall targeting antibiotics, such as *β*-lactams that disrupt the stability of the peptidoglycan meshwork are translocated by OmpF porins to induce envelope-stress response (CpX) [47]. Activation of CpX system decreases *ompF* expression [48], creating a negative feedback loop (Fig. 1B), resulting in lower porin number and lower membrane permeability (*P*_in_). Similarly, when *E. coli* are exposed to DNA targeting antibiotics that are also translocated inside the cell by OmpF, expression level of *ompF* decreases within 30-120 min after antibiotic treatment [45]. In addition to controlling antibiotic influx, bacteria can decrease intercellular antibiotic concentration through over-expression of efflux pumps [45, 49, 50] (Fig. 1B). A reduction in porin number will act in synergy with a reduction of *S/V* to confer stronger resistant phenotypes. In the future, time-lapse experiments are necessary to reveal the timescales associated with the onset and completion of morphological transformation under antibiotic perturbations, and how these timescales compare with changes in protein expression profiles responsible for regulating antibiotic influx and efflux. These studies would be essential to quantify contributions of the different resistance pathways and their synergistic effects responsible for increasing bacterial fitness.

## Acknowledgements

We gratefully acknowledge funding from EPSRC (EP/R029822/1), Royal Society (URF/R1/180187, RGF/EA/181044), and the NIH (R35 GM143042). We thank Javier López Garrido for useful comments on the manuscript.

## Author Contributions

NO and SB conceived and designed the research. NO, DS and SB developed the quantitative models. NO and DS performed simulations and data analysis. NO and SB wrote the paper.

## Notes

### Competing Interest Statement

The authors have declared no competing interest.

## References

[1] WHO report (2019).

[2] M. A. Kohanski, D. J. Dwyer, and J. J. Collins, Nature Reviews Microbiology 8, 423 (2010).

[3] F. Baquero and B. R. Levin, Nature Reviews Microbiology, 1 (2020).

[4] J. M. Blair, M. A. Webber, A. J. Baylay, D. O. Ogbolu, and L. J. Piddock, Nature Reviews Microbiology 13, 42 (2015).

[5] J. Anes, M. P. McCusker, S. Fanning, and M. Martins, Frontiers in microbiology 6, 587 (2015).

[6] S. Dam, J.-M. Pages, and M. Masi, Microbiology 164, 260 (2018).

[7] A. H. Delcour, Biochimica et Biophysica Acta (BBA)-Proteins and Proteomics 1794, 808 (2009).

[8] J. M. Blair, G. E. Richmond, and L. J. Piddock, Future microbiology 9, 1165 (2014).

[9] A. A. M. Al Mamun, M.-J. Lombardo, C. Shee, A. M. Lisewski, C. Gonzalez, D. Lin, R. B. Nehring, C. Saint-Ruf, J. L. Gibson, R. L. Frisch, et al., Science 338, 1344 (2012).

[10] K. Bush and P. A. Bradford, Nature Reviews Microbiology 17, 295 (2019).

[11] N. Ojkic, E. Lilja, S. Direito, A. Dawson, R. J. Allen, and B. Waclaw, Antimicrobial Agents and Chemotherapy 64 (2020).

[12] Z. Yao, D. Kahne, and R. Kishony, Molecular Cell 48, 705 (2012).

[13] P. Nonejuie, M. Burkart, K. Pogliano, and J. Pogliano, Proceedings of the National Academy of Sciences 110, 201311066 (2013).

[14] L. G. Monahan, L. Turnbull, S. R. Osvath, D. Birch, I. G. Charles, and C. B. Whitchurch, Antimicrobial Agents and Chemotherapy 58, 1956 (2014).

[15] L. K. Harris and J. A. Theriot, Cell 165, 1479 (2016).

[16] K. M. Mickiewicz, Y. Kawai, L. Drage, M. C. Gomes, F. Davison, R. Pickard, J. Hall, S. Mostowy, P. D. Aldridge, and J. Errington, Nature communications 10, 1 (2019).

[17] N. Ojkic, D. Serbanescu, and S. Banerjee, Elife 8, e47033 (2019).

[18] S. Banerjee, K. Lo, N. Ojkic, R. Stephens, N. F. Scherer, and A. R. Dinner, Nature Physics 17, 403 (2021).

[19] K. D. Young, Microbiology and Molecular Biology Reviews 70, 660 (2006).

[20] A. Lamsa, J. Lopez-Garrido, D. Quach, E. P. Riley, J. Pogliano, and K. Pogliano, ACS chemical biology 11, 2222 (2016).

[21] H. H. Htoo, L. Brumage, V. Chaikeeratisak, H. Tsunemoto, J. Sugie, C. Tribuddharat, J. Pogliano, and P. Nonejuie, Antimicrobial agents and chemotherapy 63 (2019).

[22] H. Shi, Y. Hu, P. D. Odermatt, C. G. Gonzalez, L. Zhang, J. E. Elias, F. Chang, and K. C. Huang, Nature Communications 12, 1 (2021).

[23] S. Vadia, L. T. Jessica, R. Lucena, Z. Yang, D. R. Kellogg, J. D. Wang, and P. A. Levin, Current Biology 27, 1757 (2017).

[24] J. López-Garrido, N. Ojkic, K. Khanna, F. R. Wagner, E. Villa, R. G. Endres, and K. Pogliano, Cell 172, 758 (2018).

[25] L. Pasquina-Lemonche, J. Burns, R. Turner, S. Kumar, R. Tank, N. Mullin, J. Wilson, B. Chakrabarti, P. Bul-lough, S. Foster, et al., Nature 582, 294 (2020).

[26] J. Elf, K. Nilsson, T. Tenson, and M. Ehrenberg, Physical Review Letters 97, 258104 (2006).

[27] D. Fange, K. Nilsson, T. Tenson, and M. Ehrenberg, Proceedings of the National Academy of Sciences 106, 8215 (2009).

[28] S. A. Angermayr, G. Chevereau, and T. Bollenbach, bioRxiv, 2020 (2021).

[29] C. Billaudeau, A. Chastanet, Z. Yao, C. Cornilleau, N. Mirouze, V. Fromion, and R. Carballido-López, Nature communications 8, 1 (2017).

[30] L. K. Harris and J. A. Theriot, Trends in Microbiology (2018).

[31] M. F. Dion, M. Kapoor, Y. Sun, S. Wilson, J. Ryan, A. Vigouroux, S. van Teeffelen, R. Oldenbourg, and E. C. Garner, Nature microbiology, 1 (2019).

[32] J. T. Sauls, S. E. Cox, Q. Do, V. Castillo, Z. Ghulam-Jelani, and S. Jun, Mbio 10 (2019).

[33] O. M. Romano and M. C. Lagomarsino, Physical Review E 101, 042403 (2020).

[34] M. Panlilio, J. Grilli, G. Tallarico, I. Iuliani, B. Sclavi, P. Cicuta, and M. C. Lagomarsino, Proceedings of the National Academy of Sciences 118 (2021).

[35] A. Typas, M. Banzhaf, C. A. Gross, and W. Vollmer, Nature Reviews Microbiology 10, 123 (2012).

[36] H. Jiang and S. X. Sun, Physical review letters 105, 028101 (2010).

[37] E. R. Rojas, K. C. Huang, and J. A. Theriot, Cell Systems 5, 578 (2017).

[38] E. R. Oldewurtel, Y. Kitahara, and S. van Teeffelen, Proceedings of the National Academy of Sciences 118 (2021).

[39] F. Si, D. Li, S. E. Cox, J. T. Sauls, O. Azizi, C. Sou, A. B. Schwartz, M. J. Erickstad, Y. Jun, X. Li, et al., Current Biology 27, 1278 (2017).

[40] P. Greulich, M. Scott, M. R. Evans, and R. J. Allen, Molecular systems biology 11, 796 (2015).

[41] M. Scott, C. W. Gunderson, E. M. Mateescu, Z. Zhang, and T. Hwa, Science 330, 1099 (2010).

[42] D. Serbanescu, N. Ojkic, and S. Banerjee, Cell Reports 32, 108183 (2020).

[43] J. Cama, M. Voliotis, J. Metz, A. Smith, J. Iannucci, U. F. Keyser, K. Tsaneva-Atanasova, and S. Pagliara, Lab on a Chip 20, 2765 (2020).

[44] A. J. Lopatkin, S. C. Bening, A. L. Manson, J. M. Stokes, M. A. Kohanski, A. H. Badran, A. M. Earl, N. J. Cheney, J. H. Yang, and J. J. Collins, Science 371 (2021).

[45] S. Suzuki, T. Horinouchi, and C. Furusawa, Nature communications 5, 1 (2014).

[46] H.-K. Ropponen, R. Richter, A. K. Hirsch, and C.-M. Lehr, Advanced Drug Delivery Reviews (2021).

[47] A. Delhaye, J.-F. Collet, and G. Laloux, MBio 7 (2016).

[48] E. Batchelor, D. Walthers, L. J. Kenney, and M. Goulian, Journal of bacteriology 187, 5723 (2005).

[49] D. Du, X. Wang-Kan, A. Neuberger, H. W. van Veen, K. M. Pos, L. J. Piddock, and B. F. Luisi, Nature Reviews Microbiology 16, 523 (2018).

[50] I. El Meouche and M. J. Dunlop, Science 362, 686 (2018).

