## Supplemental material for "Antibiotic resistance via bacterial cell shape-shifting"

**N. Ojkic, D. Serbanescu and S. Banerjee**

### Mathematical model for intracellular antibiotic dynamics

Dynamics of antibiotic concentration inside of the cell, $a_{\mathrm{in}}$, and antibiotic-substrate concentration, $x$, are given by [1]:

$$\frac{\mathrm{d}a_{\mathrm{in}}}{\mathrm{d}t} = P_{\mathrm{in}}(a_{\mathrm{out}} - a_{\mathrm{in}})\frac{S}{V} - P_{\mathrm{out}}a_{\mathrm{in}}\frac{S}{V} - k_{\mathrm{on}}a_{\mathrm{in}}x + k_{\mathrm{off}}(x_0 - x) - ka_{\mathrm{in}}, \tag{1}$$

$$\frac{\mathrm{d}x}{\mathrm{d}t} = k_x + k_{\mathrm{off}}(x_0 - x) - k_{\mathrm{on}}a_{\mathrm{in}}x - kx, \tag{2}$$

where, $a_{\mathrm{in}}$ is the external concentration of antibiotic, $x_0$ is the substrate concentration without antibiotic, $k_{\mathrm{on}}$ is the antibiotic binding rate to the substrate, $k_{\mathrm{off}}$ is the antibiotic off rate that we will assume is negligible without loosing any generality, $P_{\mathrm{in}}$ and $P_{\mathrm{out}}$ are the membrane permeability coefficients in inward and outward directions respectively. The last term on the right hand side of Eq. (1) and (2) is the antibiotic dilution term due to cell growth, where $k$ is the bacterial growth rate. Numerical solution of the above system of equations for different values of $S/V$ predicts antibiotic dynamics inside of the cell over time (Fig. 3A). We can solve the above equations analytically in two different limits to illustrate the contribution of $S/V$ transformation in reducing the intracellular antibiotic concentration.

**Weak antibiotic binding.** In the case of weak antibiotic binding, the substrate concentration $x$ is not changing significantly. Eq. (1) then becomes:

$$\frac{\mathrm{d}a_{\mathrm{in}}}{\mathrm{d}t} = P_{\mathrm{in}}a_{\mathrm{out}}\frac{S}{V} - (P_{\mathrm{in}}\frac{S}{V} + P_{\mathrm{out}}\frac{S}{V} + k')a_{\mathrm{in}}, \tag{3}$$

where $k' = k + k_{\mathrm{on}}x$. The solution of the above equation is given by:

$$a_{\mathrm{in}}(t) = a_{\mathrm{out}}\frac{P_{\mathrm{in}}\frac{S}{V}}{(P_{\mathrm{in}} + P_{\mathrm{out}})\frac{S}{V} + k'}(1 - \mathrm{e}^{-((P_{\mathrm{in}}+P_{\mathrm{out}})\frac{S}{V}+k')t}). \tag{4}$$

For the passively translocated antibiotics, equilibrated internal concentration is always smaller than the external concentration since $P_{\mathrm{in}}\frac{S}{V} < (P_{\mathrm{in}} + P_{\mathrm{out}})\frac{S}{V} + k'$. At steady-state, the intracellular antibiotic concentration is given by:

$$a_{\mathrm{in}} = a_{\mathrm{out}}\frac{P_{\mathrm{in}}\frac{S}{V}}{(P_{\mathrm{in}} + P_{\mathrm{out}})\frac{S}{V} + k}. \tag{5}$$

**Strong antibiotic binding.** In the case of strong antibiotic binding ($k_{\mathrm{on}} \gg k_{\mathrm{off}}$), we can simultaneously solve Eq. (1) and Eq. (2) to obtain the steady-state antibiotic concentration:

$$a_{\mathrm{in}} = \frac{\sqrt{k_0^2 + 4a_{\mathrm{out}}kk_{\mathrm{on}}P_{\mathrm{in}}\frac{S}{V}(k + (P_{\mathrm{in}} + P_{\mathrm{out}})\frac{S}{V})} - k_0}{2k_{\mathrm{on}}((P_{\mathrm{in}} + P_{\mathrm{out}})\frac{S}{V} + k)}, \tag{6}$$

where $k_0 = k^2 + k(P_{\text{in}} + P_{\text{out}})\frac{S}{V} + k_{\text{on}}(k_x - a_{\text{out}}P_{\text{in}}\frac{S}{V})$. Eq. (6) predicts how the steady-state concentration depends on $S/V$ (Fig. 3B). A simple expression for the antibiotic concentration inside of the cell is obtained in the limit of strong antibiotic binding to its target ($k_{\text{on}} \to \infty$):

$$a_{\text{in}} = \frac{a_{\text{out}}P_{\text{in}}\frac{S}{V} - k_x}{(P_{\text{in}} + P_{\text{out}})\frac{S}{V} + k} \,. \tag{7}$$

**Estimating membrane permeability.** Using Eq. (4) we can estimate $P_{\text{in}}$ / $P_{\text{out}}$ from the time-dependent accumulation of fluorescent fluoroquinolone antibiotics inside of the cell [2–4]. Fluoroquinolone antibiotics are predominantly translocated inside of the cell through OmpF porins and only a handful of fluoroquinolone molecules bind to their targets for concentrations below MIC [5]. If the antibiotic remains fluorescent in the substrate-bound form, the steady-state concentration inside of the cell is given by Eq. (5). By inspecting the growth of the cells in micro-fluidic devices [4], we observed that cells did not drastically elongate during the time course of the experiment ($\approx$ 6.7 min). Therefore the antibiotic dilution by growth is negligible ($k \to 0$ h$^{-1}$). Consequently, $P_{\text{in}}$ / $P_{\text{out}}$ is derived from Eq. (5):

$$\frac{P_{\text{in}}}{P_{\text{out}}} = \frac{a_{\text{in}}/a_{\text{out}}}{1 - a_{\text{in}}/a_{\text{out}}} \,. \tag{8}$$

**Antibiotic dilution by shape changes.** To estimate the amount of antibiotic dilution obtained via solely cell shape changes, we assume that bacteria go through a shape transformation without drastically changing other physiological parameters in response to antibiotic treatment. At steady-state, balanced biosynthesis requires that the substrate production rate is proportional to growth rate, $k_x = \alpha k$, where $\alpha$ is a constant [6]. Difference in the steady-state intracellular antibiotic concentrations for two bacterial shapes characterised by surface-to-volume ratios $(\frac{S}{V})_{\text{max}}$ and $(\frac{S}{V})_{\text{min}}$ is given by:

$$\Delta a_{\text{in}} = \frac{a_{\text{out}}P_{\text{in}}(\frac{S}{V})_{\text{max}} - \alpha k}{k + (P_{\text{in}} + P_{\text{out}})(\frac{S}{V})_{\text{max}}} - \frac{a_{\text{out}}P_{\text{in}}(\frac{S}{V})_{\text{min}} - \alpha k}{k + (P_{\text{in}} + P_{\text{out}})(\frac{S}{V})_{\text{min}}}. \tag{9}$$

Interestingly, difference in the steady state antibiotic concentration, $\Delta a_{\text{in}}$, is a non-monotonic function of $P_{\text{in}}/P_{\text{out}}$, reaching a maximum for optimal values of $P_{\text{in}}/P_{\text{out}}$ and the growth rate $k$. Maximum dilution of antibiotics by solely $S/V$ changes is obtained for $P_{\text{in}} \gg P_{\text{out}}$. In this limit, we maximized Eq. (9) with respect to the parameters $k$ and $P_{\text{in}}$ (Fig. 3C) to obtain $P_{\text{in}} = \frac{k}{\sqrt{(\frac{S}{V})_{\text{max}}(\frac{S}{V})_{\text{min}}}}$. This results in the following maximum value for the difference in antibiotic concentration:

$$\Delta a_{\text{in}}^{\text{max}} = (\alpha + a_{\text{out}})\frac{\sqrt{(\frac{S}{V})_{\text{max}}} - \sqrt{(\frac{S}{V})_{\text{min}}}}{\sqrt{(\frac{S}{V})_{\text{max}}} + \sqrt{(\frac{S}{V})_{\text{min}}}}. \tag{10}$$

In the case of slow substrate production or large antibiotic concentration ($a_{\text{out}} >> \alpha$) maximum dilution of the intercellular antibiotic concentration solely due to $S/V$ transformation becomes:

$$\delta_{\text{max}} = \frac{\Delta a_{\text{in}}^{\text{max}}}{a_{\text{out}}} = \frac{\sqrt{(\frac{S}{V})_{\text{max}}} - \sqrt{(\frac{S}{V})_{\text{min}}}}{\sqrt{(\frac{S}{V})_{\text{max}}} + \sqrt{(\frac{S}{V})_{\text{min}}}}. \tag{11}$$

### Single-cell growth and shape dynamics under ribosome-targeting antibiotics

To investigate the dynamic response of cell shape and growth to applied antibiotics we simulated single-cell growth over multiple generations as described previously in Serbanescu *et al.* [7]. Briefly, we first initiated cells at random life-cycle stages, and upon division followed the daughter cells over a number of generations until the steady-state is reached. During each cell generation we evolved the following six coupled differential equations (Eqs. 12-17) for cell volume $V$, division protein abundance $X$, surface area $S$, antibiotic concentration inside the cell $a_{\mathrm{in}}$, active ribosomes $r_{\mathrm{a}}$ and inactive antibiotic-bound ribosomes $r_{\mathrm{b}}$.

The cell volume rate and surface-area rate are proportional to the instantaneous volume $V(t)$, as proposed by Harris and Theriot [8]:

$$\frac{\mathrm{d}V}{\mathrm{d}t} = k(r_{\mathrm{a}})V(t), \tag{12}$$

$$\frac{\mathrm{d}S}{\mathrm{d}t} = \beta(r_{\mathrm{a}})V(t), \tag{13}$$

where $k(r_a)$ is the growth rate and $\beta(r_a)$ is the surface area synthesis rate. The expression for $\beta$ used in our simulations differs from that proposed in [8] and is shown below. The division proteins accumulate at a rate proportional to the cell volume and are degraded at a constant rate $\mu$:

$$\frac{\mathrm{d}X}{\mathrm{d}t} = \kappa_F(r_{\mathrm{a}})V(t) - \mu X(t). \tag{14}$$

The rate of growth ($k$), surface area synthesis ($\beta$) and division protein synthesis ($\kappa_F$) are given as a function of active ribosomes:

$$k(r_{\mathrm{a}}) = \kappa_t(r_{\mathrm{a}} - r_{\mathrm{min}}),$$

$$\beta(r_{\mathrm{a}}) = \nu\,[\kappa(r_{\mathrm{a}})]^{2/3}\,[\kappa_F(r_{\mathrm{a}})]^{1/3},$$

$$\kappa_F(r_{\mathrm{a}}) = \kappa_F^0(r^*_{\mathrm{max}} - r_{\mathrm{a}}),$$

where $\kappa_t$ is the translational capacity, $r_{\mathrm{min}}$ is the minimum ribosomal fraction needed for growth, $\kappa_F^0$ is the rate of production of division proteins per ribosomes, $r^*_{\mathrm{max}}$ is the ribosome mass fraction when the growth rate is maximum and $\nu$ is a geometric parameter: $\nu = \eta\pi\,[(\eta\pi/4) - (\pi/12)]^{-2/3}$, with $\eta$ the aspect ratio of the cell ($\eta \approx 4$ for *E. coli* cells).

The changes in intracellular antibiotic concentration, active ribosomes and inactive ribosomes are described by the following equations used in [7] that extend previous work[1, 9, 10]:

$$\frac{\mathrm{d}a_{\mathrm{in}}}{\mathrm{d}t} = -k(r_a)a_{\mathrm{in}} + f(a_{\mathrm{in}}, r_{\mathrm{a}}, r_{\mathrm{b}}) + J_a(a_{\mathrm{in}}, S, V) \tag{15}$$

$$\frac{\mathrm{d}r_{\mathrm{a}}}{\mathrm{d}t} = -k(r_{\mathrm{a}})r_{\mathrm{a}} + f(a_{\mathrm{in}}, r_{\mathrm{a}}, r_{\mathrm{b}}) + s(r_{\mathrm{a}}, \kappa_{\mathrm{specific}}) \tag{16}$$

$$\frac{\mathrm{d}r_{\mathrm{b}}}{\mathrm{d}t} = -k(r_{\mathrm{b}})r_{\mathrm{b}} - f(a_{\mathrm{in}}, r_{\mathrm{a}}, r_{\mathrm{b}}) \tag{17}$$

where the antibiotic influx is $J_{\mathrm{a}}(a_{\mathrm{in}}, S, V) = (P_{\mathrm{in}}(a_{\mathrm{out}} - a_{\mathrm{in}}) - P_{\mathrm{out}} a_{\mathrm{in}})\frac{S}{V}$ and $P_{\mathrm{in}}$ and $P_{\mathrm{out}}$ are the cell envelope permeabilities in the inward and outward directions. The remaining two terms are the ribosome-antibiotic interactions: $f(a_{\mathrm{in}}, r_{\mathrm{a}}, r_{\mathrm{b}}) = -k_{\mathrm{on}} a_{\mathrm{in}}(r_{\mathrm{a}} - r_{\mathrm{min}}) + k_{\mathrm{off}} r_{\mathrm{b}}$, where $k_{\mathrm{on}}$ is the rate of antibiotic binding to ribosomes and $k_{\mathrm{off}}$ is the unbinding rate, and a source term capturing how cells produce more ribosomes under translation inhibition to account for the inactive ribosomes: $s(r_{\mathrm{a}}, k_{\mathrm{specific}}) = k(r_{\mathrm{a}})(r_{\mathrm{max}} - k(r_{\mathrm{a}})(r_{\mathrm{max}} - r_{\mathrm{min}})(k_{\mathrm{specific}}^{-1} - 1/\kappa_t(r_{\mathrm{max}} - r_{\mathrm{min}})))$, where $k_{\mathrm{specific}}$ is the growth rate imposed by the growth medium in the absence of antibiotics.

**Relationship between cell shape, rate of growth and protein synthesis.** By solving Eqs. (12) and (14) between birth ($t = 0$) and division ($t = \tau$) in the limit when the growth rate is larger than the degradation rate (i.e. $k \gg \mu$), we obtain that the added division protein number is $\Delta X = X(t = \tau) - X(t = 0) = \frac{\kappa_F}{k}\Delta V$, where $\Delta V = V(t = \tau) - V(t = 0)$, the added volume per generation. At steady-state the newborn cell volume asymptotes to the added volume $V(t = 0) \approx \Delta V$. The average cell volume is therefore

$$V = \frac{k}{k_F}\ , \tag{18}$$

where $k_F = \frac{\kappa_F}{2\Delta X ln(2)}$. The accumulated division protein number is constant in a given nutrient condition and therefore $\Delta X$ can be absorbed together with other constants in $k_F$. For *E. coli* cells the average surface area and the volume are related by the power law $S = 2\pi V^{2/3}$ [11] which together with Eq. (18) leads to an elegant relationship between the surface to volume ratio and the rates of growth and division protein synthesis:

$$\frac{S}{V} = 2\pi \left(\frac{k_F}{k}\right)^{1/3}\ . \tag{19}$$

**Simulation protocol and the choice of parameters**. For each cell cycle, Eqs. (12)-(17) are evolved for $t < \tau$, where $\tau$ is the division time. Division is triggered when $X = X_0$ with $X_0$ constant. Upon division, parameters of the new daughter cells are calculated from the mother cell. We set $V_{\mathrm{new}}(0) = D_R V_{\mathrm{old}}(\tau)$, $X_{\mathrm{new}}(0) = 0$, $S_{\mathrm{new}}(0) = D_R S_{\mathrm{old}}(\tau)$, $a_{\mathrm{in}}^{\mathrm{new}}(0) = a_{\mathrm{in}}^{\mathrm{old}}(\tau)$, $r_{\mathrm{a}}^{\mathrm{new}}(0) = r_{\mathrm{a}}^{\mathrm{old}}(\tau)$, $r_{\mathrm{b}}^{\mathrm{new}}(0) = r_{\mathrm{b}}^{\mathrm{old}}(\tau)$, where $D_R$ is a Gaussian random variable with mean 0.5 and standard deviation 0.05. The parameter values are taken from Ref. [7].

In Fig. 4B we simulated the growth of the cells at steady state for 5 h to record the average values of volume, surface area and ribosome concentration in the absence of antibiotics and then the antibiotic perturbation is applied after 10 h from the start of the simulations and continued for another 20 h, when we compute the average values for the cellular components. The value of $P_{\text{out}}$ is fixed while the ratio $P_{\text{in}}/P_{\text{out}}$ is medium-dependent. The values used are the same as the ones identified in [7]. The binding and unbinding rates for antibiotics are $k_{\text{on}} = 10\,\mu M^{-1}\text{h}^{-1}$ and $k_{\text{off}} = 25\,\text{h}^{-1}$, same as the ones used in [7].

In Fig. 4C we simulated the growth of cells at steady state for 100 h in the absence of antibiotics followed by growth in the presence of antibiotics at $a_{\text{out}} = 5\,\mu M$ keeping constant extracellular antibiotic concentration. The antibiotic dilution is calculates as $\delta(a_{\text{in}}) = \frac{a_{\text{in}}^{\text{ss}}(S/V=15)-a_{\text{in}}^{\text{ss}}(S/V=3)}{a_{\text{out}}}$, where $a_{\text{in}}^{\text{ss}}$ is the steady state intracellular antibiotic concentration in the presence of $5\mu M$ extracellular antibiotic concentration and $(S/V = 3)$ and $(S/V = 15)$ denote a surface-to-volume ratio of $3\,\mu m^{-1}$ and $15\,\mu m^{-1}$ respectively, as in Fig. 3C-D. The unbinding antibiotic-ribosome rate $k_{\text{off}} = 25\,\text{h}^{-1}$ is kept fixed, while the binding rate changes such that we investigate a wide range for $k_{\text{on}}/k_{\text{off}}$ from 0.01 to $1000\,\mu M^{-1}$. Similarly, we fix the membrane permeability in the outward direction $P_{\text{out}} = 20\,\text{h}^{-1}\mu m^{-1}$ and vary $P_{\text{in}}$ such that the ratio $P_{\text{in}}/P_{\text{out}}$ ranges from 0.1 to 10. Other parameters used: $\kappa_n = 5\,\text{h}^{-1}$ is the nutritional capacity and $\kappa_{\text{specific}} = 2\,\text{h}^{-1}$ is the nutrient-specific growth rate in the absence of antibiotics [7].

### Supplementary References

[1] D. Fange, K. Nilsson, T. Tenson, and M. Ehrenberg, Proceedings of the National Academy of Sciences **106**, 8215 (2009).

[2] J. Vergalli, E. Dumont, J. Pajović, B. Cinquin, L. Maigre, M. Masi, M. Réfrégiers, and J.-M. Pagés, Nature protocols **13**, 1348 (2018).

[3] J. Vergalli, A. Atzori, J. Pajovic, E. Dumont, G. Malloci, M. Masi, A. V. Vargiu, M. Winterhalter, M. Réfrégiers, P. Ruggerone, *et al.*, Communications biology **3**, 1 (2020).

[4] J. Cama, M. Voliotis, J. Metz, A. Smith, J. Iannucci, U. F. Keyser, K. Tsaneva-Atanasova, and S. Pagliara, Lab on a Chip **20**, 2765 (2020).

[5] N. Ojkic, E. Lilja, S. Direito, A. Dawson, R. J. Allen, and B. Waclaw, Antimicrobial Agents and Chemotherapy **64** (2020).

[6] S. Cooper, Reviews in Cell Biology and Molecular Medicine (2006).

[7] D. Serbanescu, N. Ojkic, and S. Banerjee, Cell Reports **32**, 108183 (2020).

[8] L. K. Harris and J. A. Theriot, Cell **165**, 1479 (2016).

[9] J. Elf, K. Nilsson, T. Tenson, and M. Ehrenberg, Physical Review Letters **97**, 258104 (2006).

[10] P. Greulich, M. Scott, M. R. Evans, and R. J. Allen, Molecular systems biology **11**, 796 (2015).

[11] N. Ojkic, D. Serbanescu, and S. Banerjee, Elife **8**, e47033 (2019).
